# Ganglioside Lipids Inhibit the Aggregation of the Alzheimer’s Related Peptide Amyloid-*β*

**DOI:** 10.1101/2023.09.10.556751

**Authors:** Zenon Toprakcioglu, Akhila K. Jayaram, Tuomas P. J. Knowles

## Abstract

The aggregation of the amyloid-*β* (A*β*) peptides (A*β*42/A*β*40) into toxic amyloid fibrils and plaques is one of the molecular hallmarks in dementia and Alzheimers disease (AD). While the molecular mechanisms behind this aggregation process are not fully known, it has been shown that some biomolecules can accelerate this process while others can inhibit amyloid formation. Lipids, which are ubiquitously found in cell membranes, play a pivotal role in protein aggregation. Here, we investigate how ganglioside lipids, which are abundant in the brain and in neurons, can influence the aggregation kinetics of both A*β*42 and A*β*40. We find that ganglioside lipids can drastically inhibit the aggregation of A*β*42, while in the case of the smaller peptide (A*β*40), gangliosides can completely inhibit the aggregation process. Moreover, through kinetic analysis we show that the primary nucleation rate is greatly affected by the addition of gangliosides, and that the presence of these lipids can inhibit the primary nucleation rate of A*β*42 by 3 orders of magnitude. By means of viability assays of neuroblastoma cells (SH-SY5Y), we further demonstrate that amyloid fibrils formed in the absence of gangliosides are more toxic to these cells than amyloid fibrils formed in the presence of gangliosides, elucidating the inhibitory and potentially protective role that these lipids can play. Additionally, we show that monomeric A*β*40/A*β*42 form complexes with gangliosides, but not with other lipids such as POPS, suggesting that formation of ganglioside-A*β* complexes can act as a potential pathway towards inhibiting amyloid-*β* aggregation. Taken together, our results provide a quantitative description of how lipid molecules such as gangliosides can inhibit the aggregation of A*β* and shed light on the key factors that control these processes, especially in view of the fact that declining levels of gangliosides in neurons have been associated with ageing.

## Introduction

Alzheimers disease (AD) is the most common and prevalent neurodegenerative disease which accounts for 60-70% of all dementia cases.^1–3^ AD has been associated with the self-assembly and misfolding of physiologically generated proteins such as tau and amyloid-*β* (A*β*).^4–8^ The two major fragments of the A*β* peptide, A*β*42 and A*β*40, have been known to aggregate into fibrils, and ultimately amyloid plaques - a process which is regarded as the hallmark for AD.^7–9^ Furthermore, the aggregation of A*β* into such structures results in neuronal dysfunction and ultimately cell death, which subsequently leads to the pathology and progression of AD.^10, 11^ Since the A*β* peptide is naturally found extracellularly, due to being cleaved-off from the membrane-bound amyloid precursor protein (APP),^2, 11^ understanding the interaction of A*β* with lipid membranes and surfaces from a mechanistic viewpoint is extremely important. While it is known that different lipids can affect protein aggregation,^12, 13^ either by inhibiting or accelerating the overall process, the physicochemical properties of these lipids determine which of these pathways the reaction will undertake.

A particular type of lipid which is abundant in the brain and in neuronal cells is ganglioside. These glycosphingolipids are essential in cell-to-cell communication, cell signaling and also play an important role in immunomodulation. Additionally, they are thought to be involved with neuroprotection, have neurotrophic properties^14, 15^ and exhibit neurorestorative effects.^16, 17^ Following their synthesis, gangliosides are transported to the outer leaflet where they form part of the plasma membrane and are almost exclusively found there.^18, 19^ Their large hydrophilic head group protrudes into the extracellular environment, where they can interact with other biomolecules (proteins and/or lipids). ^20^ Gangliosides are thus physiologically expressed in the outer layer of the membrane surface and given that A*β* is also physiologically present in the extracellular space, the interaction between A*β* and ganglioside lipids is extremely important to study and understand. Although gangliosides are abundant in neurons, with concentrations reaching ten times higher than those found in non-neuronal cells, ganglioside levels progressively decrease with age.^21–24^ This could therefore be a contributing factor as to why elderly people, who naturally possess lower ganglioside levels, are more prone to neurodegeneration and dementia. Numerous studies involving gangliosides and *α*-synuclein, a protein implicated and involved with Parkinson’s disease (PD) and other synucleinopathies, have been conducted.^24–27^ To date, however, there are limited studies investigating the effect that gangliosides have on inhibiting or accelerating the aggregation of A*β*, with most studies focusing on computational and simulation results.^28, 29^

Here, in order to address this and shed light on the interaction between ganglioside lipids and A*β*, we probe the aggregation kinetics of both A*β*42 and A*β*40 and find that gangliosides can delay the aggregation of A*β*42, while completely inhibiting the self-assembly of A*β*40. Through kinetic analysis we derive the microscopic rate constants behind these aggregation phenomena, and show that the rate of primary nucleation is greatly affected by the presence of gangliosides, where in the case of A*β*42 it is reduced by 3 orders of magnitude. In comparison, we show that addition of lipids such as POPS, do not affect the primary nucleation rate of A*β*42 to the same degree as gangliosides. Furthermore, we perform viability assays of neuroblastoma cells and show that fibrils formed in the presence of gangliosides are less cytotoxic than fibrils formed in the absence of the lipid. Finally, we establish that gangliosides are able to form complexes with A*β*40/A*β*42, which was not the case for the other lipid tested (POPS), suggesting that such ganglioside-A*β* complexes may be key factors in the overall inhibition of amyloid-*β* aggregation. These results demonstrate the inhibitory and protective role that gangliosides can play in the aggregation of A*β*. More importantly, our results pave the way for new insights into the molecular mechanisms behind dementia and AD, and highlight how certain lipids can potentially be used for future healthcare applications.

## Results and Discussion

### Ganglioside Lipids Inhibit Amyloid-***β*** Aggregation

A general schematic illustrating how ganglioside lipids inhibit A*β* aggregation is shown in Scheme 1. The proposed mechanism which we describe involves an A*β*-ganglioside complex being formed which inhibits the aggregation of amyloid-*β* by decreasing the primary nucleation rate. The experiments leading to the findings supporting this proposed mechanism are described in the following sections.

**Scheme 1:**
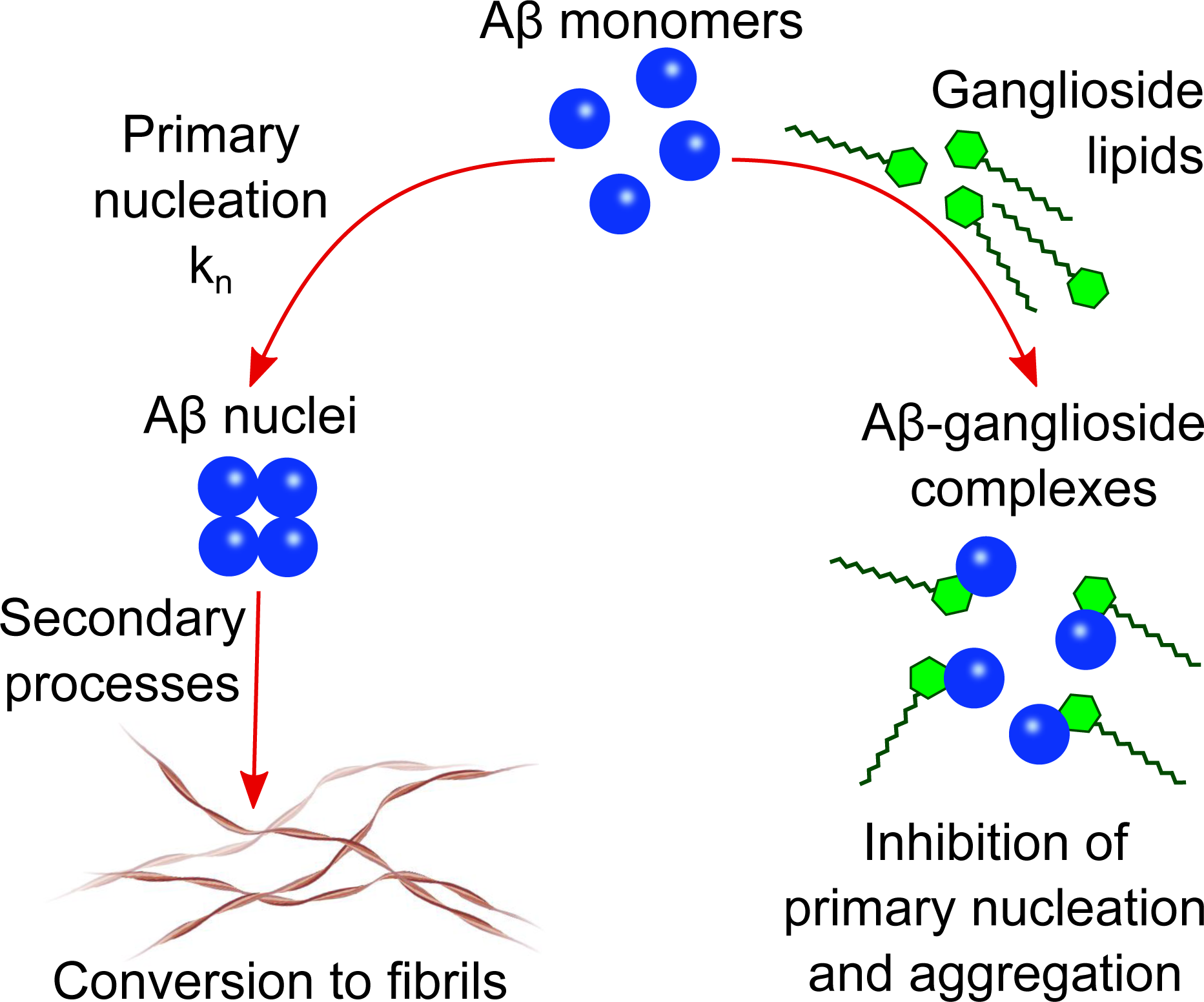
Schematic illustration of the proposed mechanism of A*β* inhibition in the presence of ganglioside lipids. A*β* forms a complex with gangliosides which in turn inhibits primary nucleation and subsequently aggregation of the peptide into amyloid fibrils.

In order to investigate how interfaces and in particular gangliosides can affect amyloid- *β* (A*β*) aggregation, we varied the nature of the interface. This was done by either leaving the protein solution within the 96-well plate as is, or by adding a ganglioside lipid solution on top of each protein solution in the 96-well plate (Figure 1a-b). This ensured that the ganglioside solution fully covered the protein phase and thus eliminated any other interfacial interaction. The kinetics of amyloid formation were then monitored by observing the increase in fluorescence of Thioflavin T (ThT). This fluorophore has the characteristic of increasing its quantum yield when bound to *β*-sheet rich structures, and therefore an increase in fluorescence intensity correlates directly with an increase in aggregated structures.

**Figure 1:**
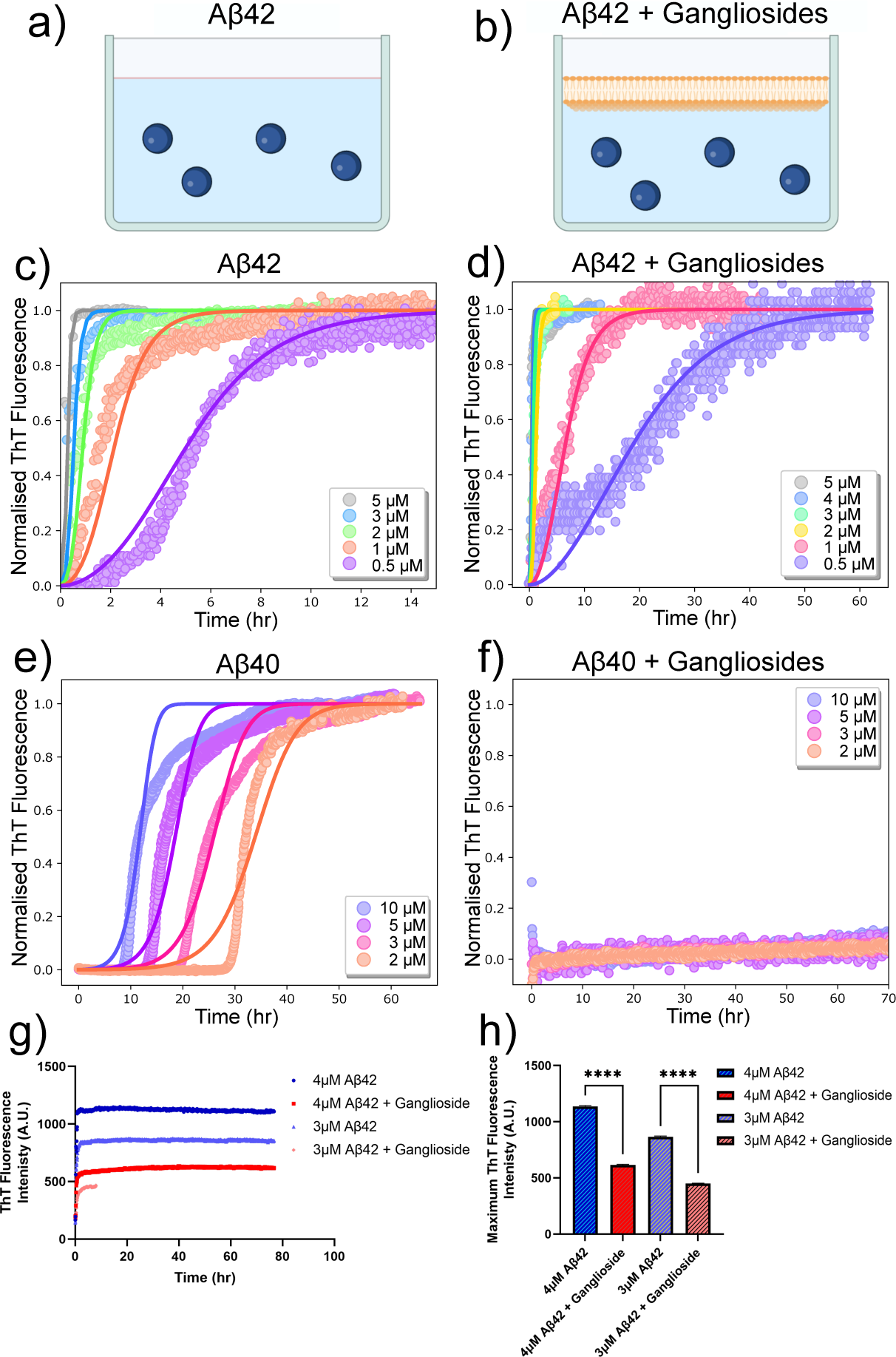
**(a-b)** Schematic showing the experimental set-up used to probe the effect of ganglioside lipids on the aggregation of the Alzheimer’s related peptide amyloid-*β* (A*β*). **(c-d)** Normalised ThT aggregation kinetics curves for A*β*42 in the absence and presence of gangliosides. Points correspond to experimental data while the solid lines correspond to the fits. **(e-f)** Normalised ThT aggregation kinetics curves for amyloid-*β* 40 (A*β*40) in the absence and presence of gangliosides. Total inhibition of A*β*40 aggregation was observed for samples containing a ganglioside interface. Points correspond to experimental data while the solid lines correspond to the fits. **(g)** Raw ThT aggregation kinetics for A*β*42 (3 and 4 *µ*M solutions) in the absence and presence of gangliosides. **(h)** Maximum ThT fluorescence intensity values obtained from the corresponding curves in (g). Data are shown as mean SD. One-way ANOVA with Tukeys multiple comparisons test.

The resulting kinetic curves (which have a characteristic sigmoidal curve) show the build-up of fibril mass over time. In these experiments, a concentration range from (0.5 *µ*M to 5 *µ*M) was investigated. It is clear from the results in Figure 1c-d, that in the absence of gangliosides, i.e. when amyloid*β* 42 (A*β*42) is left to aggregate on its own, faster aggregation kinetics are observed (Figure 1c) than when gangliosides are added to the top of the aggregation assay (Figure 1d). This inhibitory effect can be easier seen for the lowest concentration protein solutions (0.5 *µ*M), where the half-time increases from around 6 hours (Figure 1c) to 30 hours (Figure 1d) upon addition of gangliosides. Moreover, we quantified these kinetic curves by employing the use of a chemical kinetics framework which allowed us to interpret the aggregation profiles in terms of the rate constants of the underlying microscopic steps of aggregation. Such steps include primary nucleation, fibril elongation, and secondary processes such as fragmentation and surface catalysed secondary nucleation. The rate constants which correspond to these microscopic steps are k*_n_*, k_+_, and k_2_ which are the primary nucleation rate, the elongation rate, and the secondary nucleation rate constant, respectively. n*_c_* corresponds to the reaction order of the primary process and n_2_ is the reaction order of the secondary pathway. It is known that A*β* aggregates via secondary nucleation.^30, 31^ Therefore, a secondary nucleation dominated model was used to fit the data for protein samples both in the absence and presence of the ganglioside interface. It was found that in the absence of gangliosides the elongation and secondary nucleation product, k_+_k_2_, obtained through this analysis was k_+_k_2_ = 1.20×10^19^ M*^−^*^3^ h*^−^*^2^, while the elongation and primary nucleation product was found to be k_+_k*_n_* = 4.32×10^11^ M*^−^*^2^ h*^−^*^2^. Conversely, in the presence of a ganglioside interface, the corresponding combined rate constants were k_+_k_2_ = 1.23×10^17^ M*^−^*^3^ h*^−^*^2^ and k_+_k*_n_* = 2.03×10^10^ M*^−^*^2^ h*^−^*^2^. It is clear from these results that the aggregation kinetics of A*β*42 proceed much faster in the absence of gangliosides.

Moreover, a similar assay was conducted for the smaller amyloid peptide, amyloid-*β* 40 (A*β*40). It was found that in the absence of gangliosides, the peptide aggregated via a secondary nucleation pathway, as expected,^31^ with rate constants (k_+_k_2_ = 5.86×10^14^ M*^−^*^3^ h*^−^*^2^ and k_+_k*_n_* = 1.57×10^6^ M*^−^*^2^ h*^−^*^2^). The corresponding aggregation curves are shown in Figure 1e. However, upon addition of gangliosides, total inhibition of A*β*40 aggregation was observed across the same concentration range (2 - 10 *µ*M), showing the potent ability of this lipid to delay and even stop the self-assembly of A*β*40. These results are shown in Figure 1f. Furthermore, the un-normalised aggregation curves of A*β*42 for a couple of concentrations with and without gangliosides were plotted. The ThT fluorescence intensity gives us an indication of the amount of fibrils present in solution. For all concentrations tested, it was found that the solutions containing gangliosides always exhibited a decreased fluorescence signal, approximately by a factor of 2, suggesting the presence of fewer fibrils in these solutions. These data are summarised in Figure 1g-h.

### Surface-to-Volume Ratio Affects Amyloid-***β*** Aggregation by Modulating the Rate of Heterogeneous Primary Nucleation

In order to determine the effect that the surface can have on the rate of heterogeneous primary nucleation, we first investigated the effect of changing the surface-to-volume (S/V) ratio, as is shown in the schematic in Figure 2a. Kinetic experiments were thus conducted in systems with different volumes, where wells were filled with a range of 80 - 140 *µ*L.^32^ This corresponded to a S/V ratio in the range of 0.218 to 0.132 mm*^−^*^1^. The results show that for the range of S/V ratios explored, a higher S/V ratio results in faster aggregation kinetics of A*β*42, Figure 2d. Since all parameters (other than the S/V ratio) remain constant within the solution, this effect can only be attributed to an increase in the primary nucleation rate. It should also be noted that due to the fact that a ganglioside interface completely inhibited A*β*40 aggregation (Figure 1f), we chose to focus the S/V study primarily on the A*β*42 peptide.

**Figure 2:**
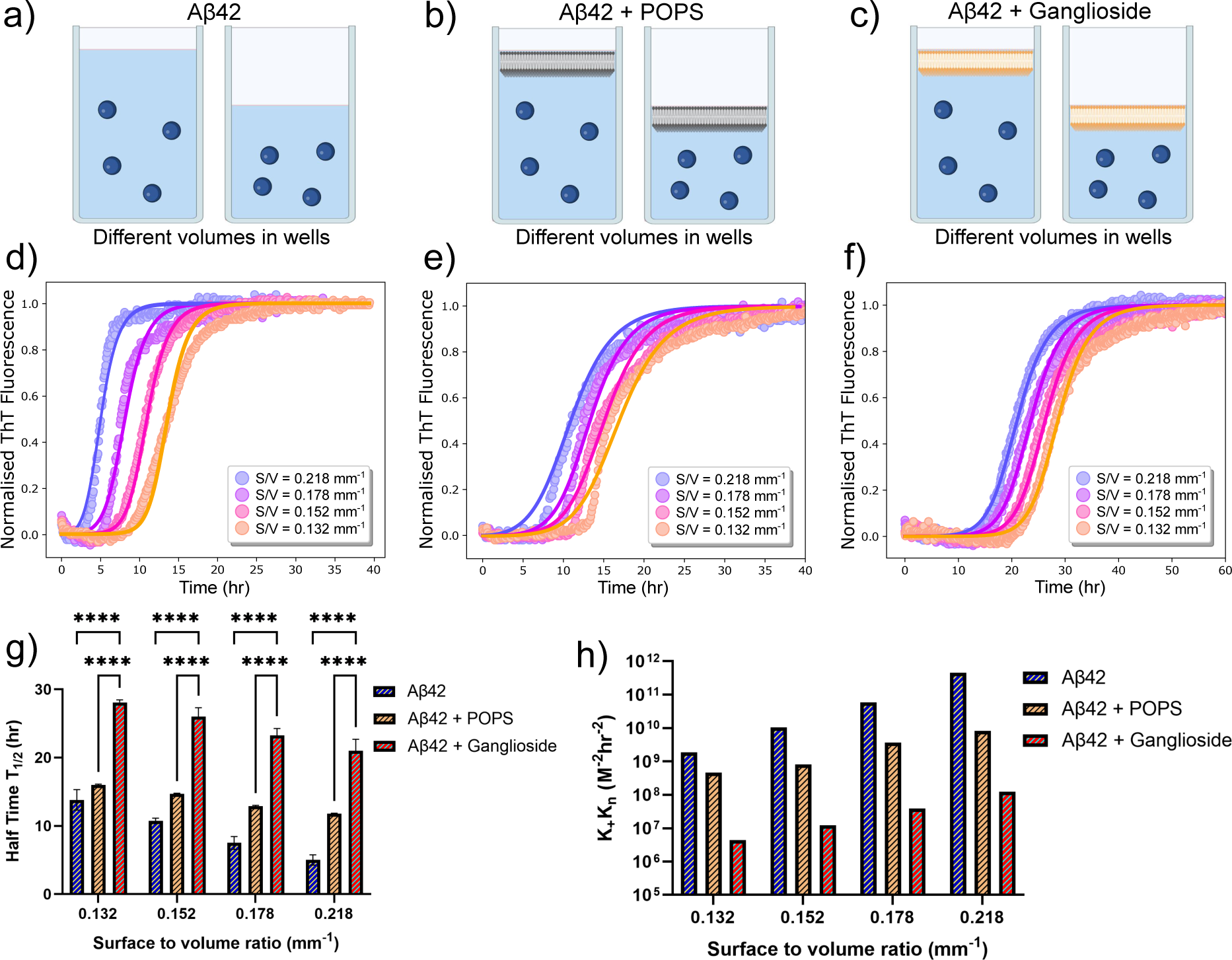
**(a)** Schematic showing how the surface-to-volume ratio (S/V) was changed in order to probe how surfaces affect heterogeneous primary nucleation. **(b-c)** Similar schematic to (a) with the addition of a POPS (b) or a ganglioside (c) interface at the top of the protein solution. **(d-f)** Normalised ThT aggregation kinetics of a 0.5 *µ*M A*β*42 solution alone (d) in the presence of POPS (e) and in the presence of gangliosides (f), for different S/V ratios. Points correspond to experimental data while the solid lines correspond to the fits. **(g)** Plot of half-time of the aggregation kinetics of a 0.5 *µ*M solution of A*β*42 as a function of the S/V ratio for all three systems (alone, with POPS and with gangliosides). Data are shown as mean SD. Two-way ANOVA with Sidak’s multiple comparisons test. ****p*<*0.0001 **(h)** Plot of k_+_k*_n_* as a function of the surface-to-volume ratio for A*β*42 with and without the presence of the two lipids (POPS and gangliosides).

We then proceeded to use the same experimental system with the incorporation of lipids. In addition to investigating the effect of gangliosides on the aggregation kinetics, we also chose to include another lipid, POPS, as a control experiment. The lipids were added to the top of the protein solution (Figure 2b-c) and the same volume range was explored. The kinetic graphs are shown in Figure 2e-f. The same trend was observed - higher S/V ratios result in faster aggregation kinetics. Moreover, while both lipids delayed A*β*42 aggregation, the inhibitory effect of gangliosides on the aggregation of A*β*42 was again observed and is clearly more prominent than that of POPS. The half-time of each S/V ratio was plotted, the results of which are shown in Figure 2g. For all S/V rations, there is a decrease in the half-time and furthermore, for the same S/V ratio, addition of lipid delays the aggregation of A*β*42. In particular, for samples where gangliosides are present, the half-time of A*β*42 is massively increased, up to a factor of 2 to 3.

To quantify the effect that the S/V ratio has on the primary nucleation rate, we employed the use of chemical kinetics once again. The fitting was first performed for one S/V ratio, and all parameters other than k_+_k*_n_* were kept constant thereafter. The k_+_k_2_ values used were taken from the analysis performed previously (Figure 1c-d). The primary nucleation rate for each S/V ratio was then obtained and plotted in Figure 2f. This analysis (Figure 2f) reveals that the elongation and secondary nucleation rate constants are independent of the S/V ratio and, more importantly, that the primary nucleation rate is greatly affected. In the absence of a lipid interface, the primary nucleation rate varies from 1.85×10^9^ M*^−^*^2^ h*^−^*^2^ to 4.51×10^11^ M*^−^*^2^ h*^−^*^2^. However, in the presence of a ganglioside interface, the primary nucleation rate varies from 4.43×10^6^ M*^−^*^2^ h*^−^*^2^ to 1.22×10^8^ M^−2^ h^−2^.

This tells us two things; that by changing the S/V ratio by less than a factor of 2, the primary nucleation rate constant is affected by more than 2 orders of magnitude, and for a given S/V ratio, the presence of a ganglioside interface can inhibit the primary nucleation rate by 3 orders of magnitude. Conversely, for a given S/V ratio, the presence of POPS only modulated the primary nucleation rate by 1 order of magnitude. These findings therefore suggests that different interfaces and lipids can strongly modulate the aggregation of A*β*42 through controlling the rate of primary nucleation.

We next sought to verify the conclusions derived from the kinetic analysis which suggested that interfaces modulate primary nucleation. This was tested by bypassing the primary nucleation step in the aggregation pathway, through addition of preformed seeds to the protein solutions in the 96-well plates (schematic shown in Figure 3a-b). Assuming that the formation of nuclei predominately occurs at the interface, addition of seeds to the solutions would effectively negate any surface dependence on overall fibrillar growth, as the nuclei required to initiate the reaction would already be in solution. This implies that there should be minimal to no differences in kinetic behaviour when changing the S/V ratio. We tested this hypothesis for an A*β*42 solution with and without the presence of either POPS or gangliosides. In all cases, our prediction that the seeds would circumvent primary nucleation was verified and it is clear from the aggregation kinetic curves in Figure 3d,f,h that independently of the S/V ratio, similar kinetics are observed upon addition of 10% seeds. This is in stark contrast with the behaviour observed for the aggregation kinetics in the absence of seeds, Figure 3c,e,f. Moreover, the half-time of each S/V ratio for unseeded A*β*42 aggregated in the absence and presence of either POPS or gangliosides is shown, where it is again clear that an increase in the S/V ratio enhances the kinetic process whereas addition of both lipids, but in particular ganglioside increases the half-time and therefore inhibits the overall reaction (Figure 3i). Conversely, the half-time plot of the corresponding seeded solutions is shown in Figure 3j, where it is clear that for all cases the half-time has converged to the same value.

**Figure 3:**
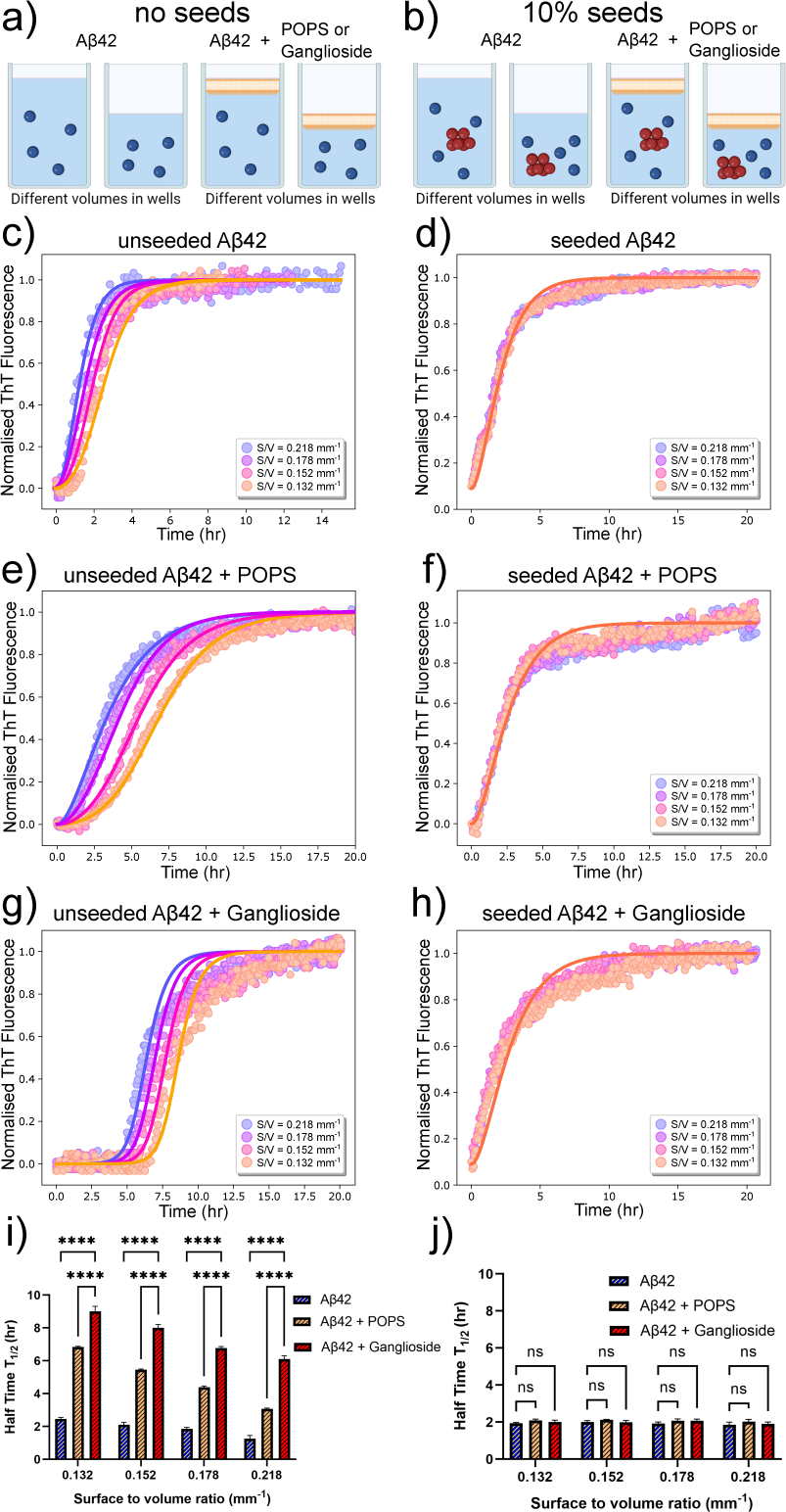
**(a-b)** Schematic showing how the surface-to-volume ratio (S/V) was changed with and without a POPS/ganglioside interface, in the presence and absence of preformed seeds. **(c)** Normalised ThT aggregation kinetics of a 1 *µ*M A*β*42 solution in the absence of gangliosides, for different S/V ratios. **(d)** Normalised ThT aggregation kinetics with the same conditions as that in (c), with the addition of 10% seeds. **(e)** Normalised ThT aggregation kinetics of a 1 *µ*M A*β*42 solution in the presence of POPS, for different S/V ratios. **(f)** Normalised ThT aggregation kinetics with the same conditions as that in (e), with the addition of 10% seeds. **(g)** Normalised ThT aggregation kinetics of a 1 *µ*M A*β*42 solution in the presence of gangliosides, for different S/V ratios. **(h)** Normalised ThT aggregation kinetics with the same conditions as that in (g), with the addition of 10% seeds. In all cases, points correspond to experimental data while the solid lines correspond to the fits. **(i)** Plot of half-time of the unseeded aggregation kinetics in (c), (e) and (g) as a function the S/V ratio. **(j)** Plot of half-time of the seeded aggregation kinetics in (d), (f) and (h) as a function the S/V ratio. Data in both (i-j) are shown as mean SD. Two-way ANOVA with Sidak’s multiple comparisons test. ****p*<*0.0001, ns = non-significant

### Morphological Changes due to the Presence of Gangliosides

As our results suggest that nucleation predominantly takes place at interfaces and that gangliosides have the ability to inhibit the aggregation of amyloid-*β* peptides, it stands to reason that the nature of the interface can modulate the morphology of the structures formed. In other words, samples with and without gangliosides could exhibit structural and morphological differences from each other. In order to explore this effect, transmission electron microscopy (TEM) was conducted on samples nucleated in the absence and presence of gangliosides and POPS. The TEM results revealed that mature fibrils were formed for both A*β*42 (Figure 4a) and A*β*40 (Figure 4g) when aggregated in the absence of any lipid. Additionally, A*β*42 and A*β*40 fibrils were also observed for the samples co-incubated with POPS (Figure 4b and Figure 4h respectively). However, in the samples where gangliosides were present, mostly oligomers and protofibrils were seen for A*β*42 (Figure 4c), and no ordered structures were observed for A*β*40 (Figure 4i).

**Figure 4:**
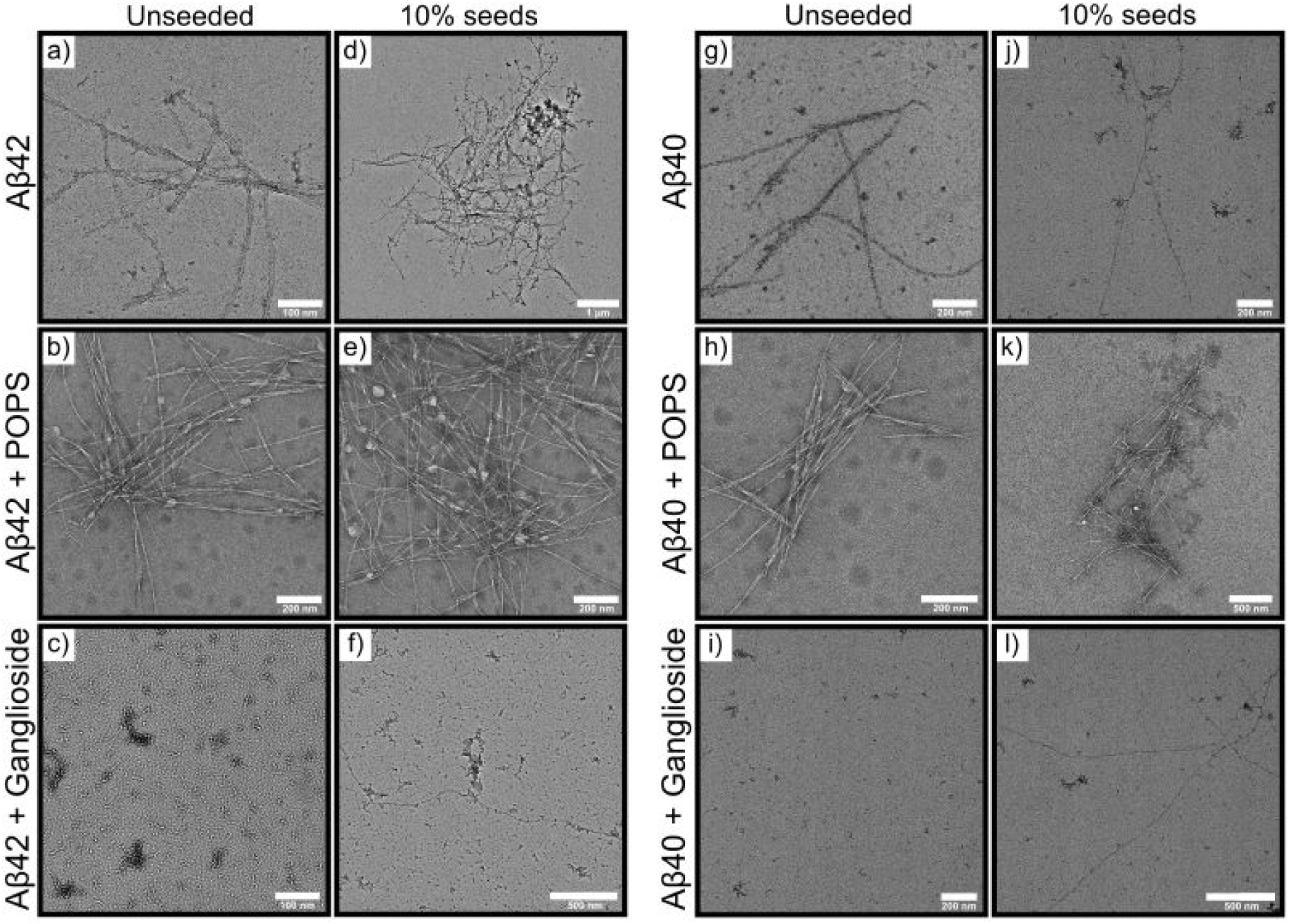
**(a-c)** TEM micrographs of A*β*42 and **(g-i)** A*β*40 following incubation of the peptides under different conditions, alone (a,g) with POPS (b,h) and in the presence of gangliosides (c,i). **(d-f)** and **(j-l)** TEM micrographs of the corresponding systems with the addition of 10% pre-formed seeds.

Moreover, TEM micrographs were acquired for both A*β*42 and A*β*40 samples co-incubated with preformed seeds. The TEM images show that under such conditions, gangliodsides have a much smaller effect on the morphology of structures formed with mature fibrils being observed for all systems (Figure 4d-f and Figure 4j-l). This is due to gangliosides having the ability to modulate the primary nucleation of the aggregation process, which is circumvented if enough seeds are added to the solution. Additionally, it should be noted that we observed fewer fibrils in seeded samples where gangliosides were present, than in the corresponding samples which did not contain any gangliosides. Conversely, multiple fibrils were found in solutions which lacked any lipids and in samples were POPS was present. These observations are in agreement with our results for unseeded data, Figure 1g-h. Combined, these findings suggest that gangliosides not only inhibit A*β* aggregation, but also reduce the overall amount of fibrils present in the solution. Furthermore, circular dichroism (CD) spectra of A*β*42 by itself, A*β*42 with POPS and A*β*42 with gangliosides were taken. The data, shown in Figure S1, suggest that A*β*42 alone (black curve) and in the presence of POPS (blue curve) form *β*-sheets. This is confirmed from the characteristic negative band at 217/218nm. However, the sample where A*β*42 was aggregated in the presence of gangliosides (red curve) displayed two peaks, one at 217/218nm and one at 207/208, suggesting that the species present in this sample are not pure aggregates, which confirms the TEM and ThT aggregation kinetic findings discussed above.

### Gangliosides Inhibit Amyloid-***β*** Aggregation within a Cell Mimicking Lipid Membrane Mixture

In order to further assess the inhibitory effect of ganglioside lipids on A*β* aggregation, we sought to mimic a cellular environment and monitor the aggregation kinetics. A mixture of different lipids, which corresponded to a typical membrane composition of a neuronal cell^33^ was prepared Figure 5a. We then sequentially doped the lipid mixture with ganglioside lipids, and monitored the aggregation kinetics for these different lipid-composition systems. It was found that in the absence of any lipid interface, A*β*42 aggregated the fastest (black points in Figure 5b). However, upon addition of the lipid mixture which mimicked the cellular environment, we can already see a delay in the aggregation kinetics. Moreover, as the amount of gangliosides are increased within the lipid mixture, the aggregation of A*β*42 is further inhibited, in a dose dependent manner, with an inhibitory plateau being reached at around 20/30% addition of gangliosides (Figure 5b). The half-time plots of the corresponding aggregation curves are shown in Figure 5c, where it is clear that the higher the amount of ganglioside within the lipid mixture, the larger the inhibitory effect. A similar trend was observed for the aggregation of A*β*40 (Figure SI S2) where in the absence of a lipid interface the peptide aggregated at a much faster rate compared to in the presence of a lipid mixture. However, upon dosing the lipid mixture with even a small amount of gangliosides (as low as 5%), total inhibition of A*β*40 aggregation was observed (Figure SI S2), which is in clear agreement with what was observed in Figure 1f.

**Figure 5:**
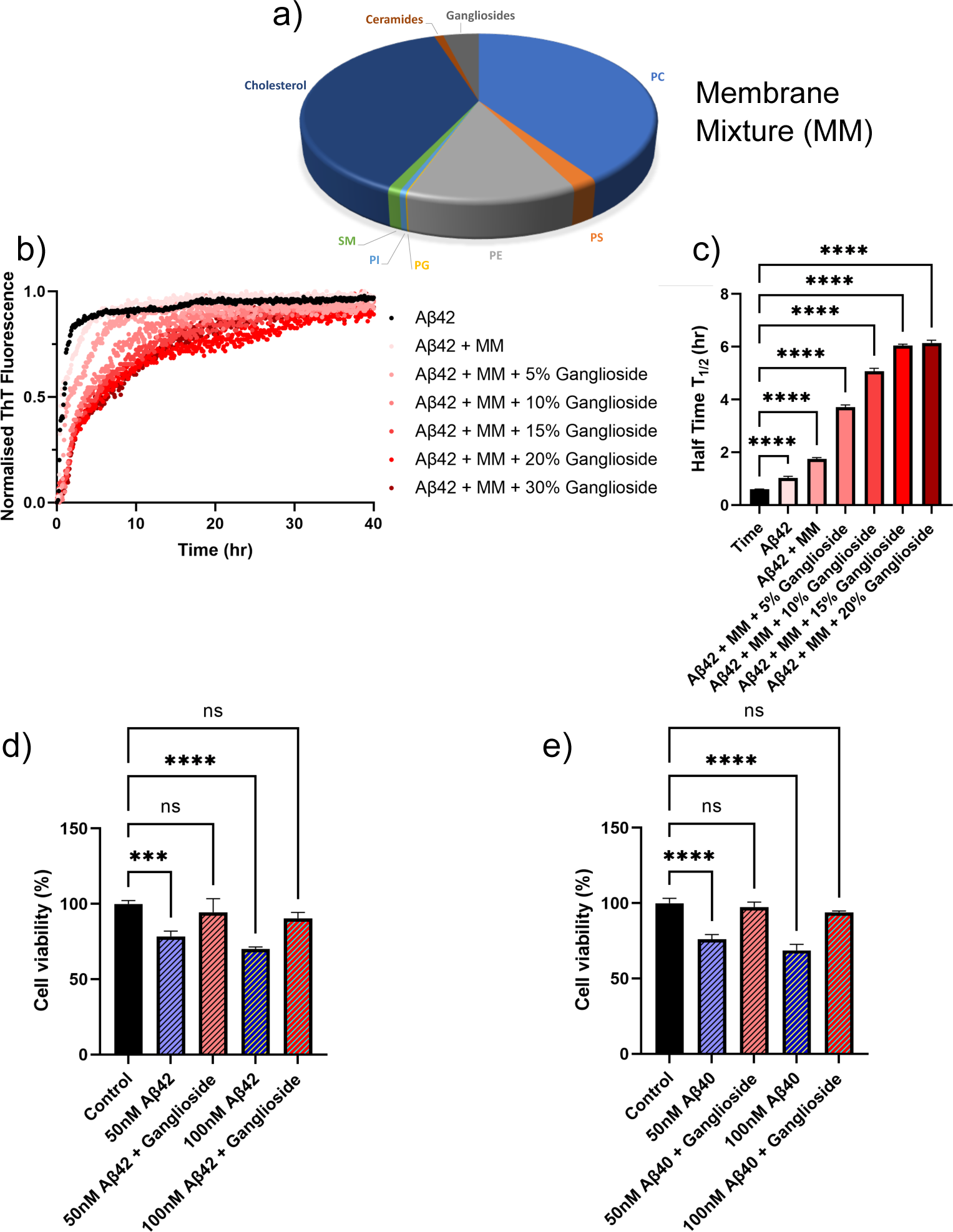
**(a)** Lipids used to mimic the membrane composition of a typical neuronal cell.^33^ **(b)** Normalised ThT aggregation kinetics of a 1 *µ*M A*β*42 solution in the presence of a lipid mixture with varying amounts of gangliosides. **(c)** Plot of half-time of the aggregation kinetics in (b) as a function of the amount of gangliosides present in the cell-mimicking lipid mixture. **(d-e)** Cytotoxicity of A*β*42 (d) and A*β*40 (e) of neuroblastoma cells (SH-SY5Y). Cell viability of SH-SY5Y cells was determined using an MTT assay following treatment with fibrils formed in the absence/presence of gangliosides. Data in both (d-e) are shown as mean SD. One-way ANOVA with with Tukeys post-hoc comparisons test. ****p*<*0.0001, ***p*<*0.001, ns = non-significant.

### The Cytotoxicity of SH-SY5Y Neuroblastoma Cells is reduced for Amyloid-***β*** Fibrils Aggregated in the Presence of Gangliosides

It is known that aggregates of A*β* possess cytotoxic properties, which have been clearly linked to neurodegeneration in Alzheimer’s Disease and other dementias. In order to obtain a more quantitative understanding as to how gangliosides may possibly protect cells against toxic amyloid fibrils, we performed assays investigating cellular viability. We assessed the cytotoxicity of fibrils formed both in the absence and presence of gangliosides with SH-SY5Y neuroblastoma cells (Figure 5d-e). It was determined that for both concentrations tested (50 and 100 nM), fibrils of A*β*42 formed in the absence of gangliosides were more cytotoxic that fibrils formed in the presence of the lipid. This can be seen Figure 5d. Moreover, in the case of A*β*40, 50 and 100 nM fibril samples were added to SH-SY5Y cells. Similarly to before, it was found that the samples where A*β*40 had been incubated in the presence of gangliosides there was minimal cytotoxicity, whereas the samples aggregated in the absence of gangliosides displayed reduced cellular viability in a dose dependent manner. These data are summarized in Figure 5e.

## Discussion

The results described in this paper provide the basis for understanding the inhibitory effect of ganglioside lipids on the aggregation of amyloid-*β* (A*β*). In the absence of a ganglioside interface, both A*β*42 and A*β*40 aggregate and self-assembly to form fibrillar structures, however, addition of gangliosides significantly inhibits the aggregation of A*β*42, while completely inhibiting the aggregation of the smaller amyloid fragment A*β*40 (Figure 1). Moreover, by varying the surface-to-volume ratio (S/V) we show that primary nucleation occurs at the interface (Figure 2a,d) and that addition of lipids at this interface not only inhibits the aggregation process (Figure 2e-f), but more importantly, modulates and decreases the primary nucleation rate. Interestingly, the lipid POPS decreases the primary nucleation rate by only one order of magnitude whereas addition of ganglioside lipids can modulate the rate by at least three orders of magnitude (Figure 2h), which in turn inhibits the overall aggregation process.

Furthermore, transmission electron microscopy revealed that the morphologies of the fibrils is greatly affected by addition of gangliosides. Both A*β*42 and A*β*40 aggregated into mature fibrils in the absence of a lipid, and in the presence of POPS. However, in the presence of gangliosides, no aggregates were observed for A*β*40 (Figure 4i), which confirms the kinetics finding in Figure 1d, while mainly protofibrillar structures were found for A*β*42 (Figure 4c). Additionally, the cellular membrane environment of a neuronal cell was mimicked by combining a mixture of different lipids, and the aggregation kinetics of A*β*42/A*β*40 in the presence of this lipid mixture was monitored. It was found that not only does a lipid mixture delay the aggregation of A*β*, but doping such a lipid mixture with gangliosides further inhibited the self-assembly of the amyloid peptides (Figure 5b-c), showcasing the importance that gangliosides present on neural membranes can play in the inhibition of A*β* aggregation. Finally, in order to assess the toxicity of aggregates formed in the absence/presence of gangliosides, cell viability assays were conducted with neuroblastoma (SH-SY5Y) cells. For both A*β*42 and A*β*40 it was observed that aggregates formed in the presence of gangliosides exhibited reduced toxicity to SH-SY5Y cells compared to fibrils formed in the absence of the lipid (Figure 5d-e). This is in direct agreement with reports of the neuroprotective effects of gangliosides with cell cultures,^34^ rat models of Alzheimer’s Diseases (AD) ^35–37^ and even results from clinical trials.^35^

The fact that a lipid interface, and in particular gangliosides which are abundant in neuronal cells, can delay amyloid formation via the inhibition of primary nucleation, the fundamental first step in the aggregation process of A*β*, is of particular interest especially in light of the fact that ganglioside abundance may decrease with ageing. ^22–24^ A schematic representation showing the overall process is shown in Figure 6. Moreover, given both the results obtained from our kinetic analysis and the cellular viability assays, what we can potentially conclude from our data is that gangliosides have a protective role against both the aggregation of A*β* peptides but also against cytotoxicity towards neuroblastoma cells (schematically represented in Figure 6d).

**Figure 6:**
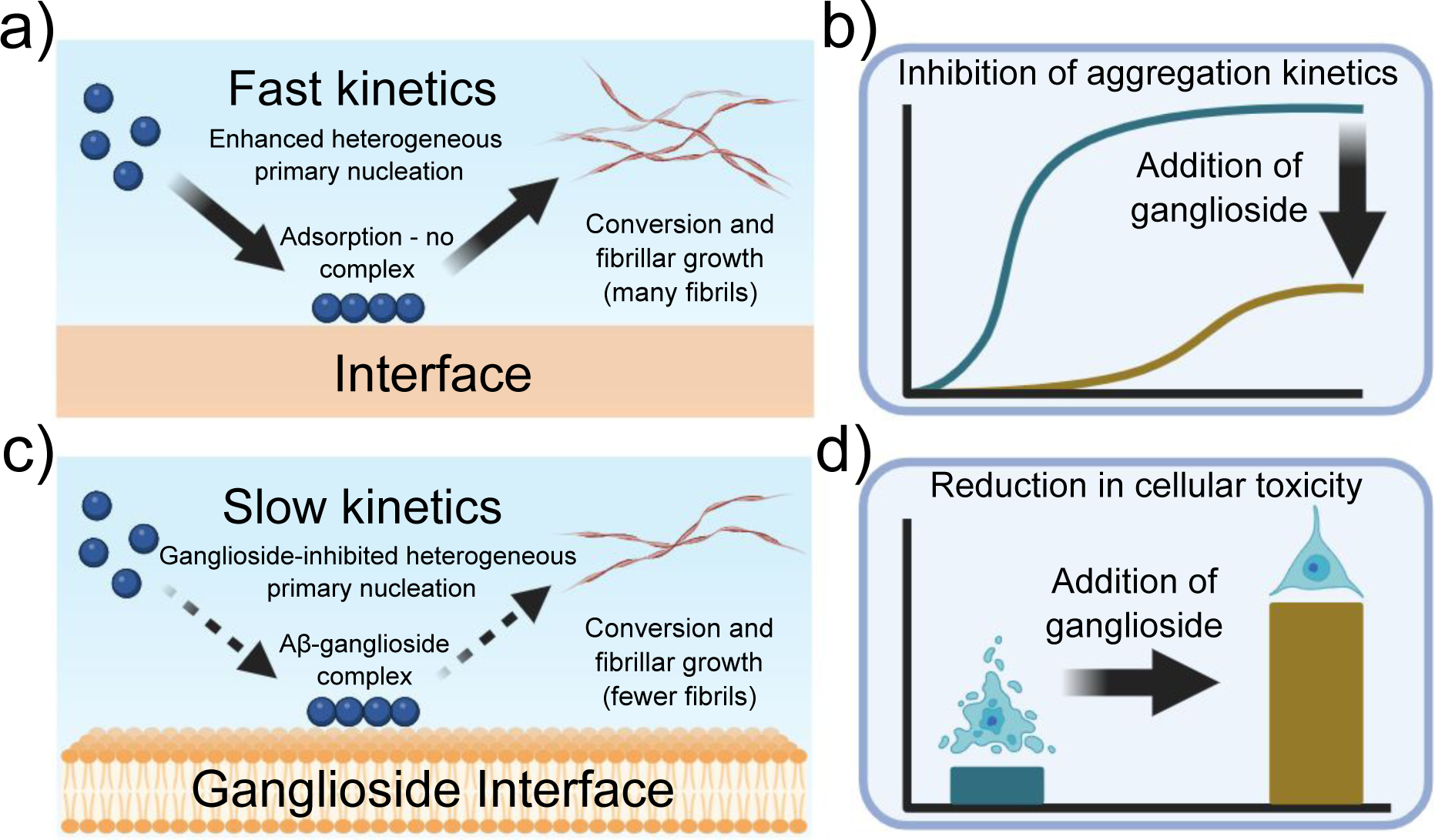
Schematic representation of the experimental findings described throughout this paper. **(a)** Primary nucleation and conversion of fibrils is enhanced in the presence of an interface, **(c)** while it is significantly inhibited when gangliosides are added, possibly due to the formation of an A*β*-ganglioside complex. **(b)** Aggregation kinetics of A*β* are inhibited in the presence of gangliosides, **(d)** while addition of lipids to the protein mixture before aggregation reduces cellular toxicity.

A potential mechanism behind the inhibitory effect that gangliosides have on the aggregation of A*β* can be associated with the lipid’s potential to form complexes with A*β*.^28, 29^ It is known that particular gangliosides such as GM1, which is a component of the outer layer of the plasma membrane, specifically binds *α*-synuclein (a protein associated with Parkinson’s Disease) in order to promote an *α*-helix structure over *β*- sheet, the latter of which is the structure that aggregated *α*-synuclein adopts.^36, 38, 39^ It is therefore quite plausible that gangliosides can be implicated in binding and forming complexes with other proteins and peptides.

In order to investigate whether this is the case for A*β*, we conducted microfluidic diffusional sizing (MDS) on A*β*40/A*β*42 samples both in the absence and presence of lipids. MDS works by measuring the degree of molecular diffusion across a microfluidic channel, which in turn can be used to calculate the apparent hydrodynamic radius. The hydrodynamic radius of A*β*40/A*β*42 peptides was initially measured, Figure 7b-c (blue bar). Following this, POPS lipids were added to monomeric A*β*40/A*β*42 and the radius was measured. No detectable change was seen, implying that POPS does not bind with A*β* monomers (yellow bars in Figure 7b-c). Finally, the same MDS experiment was conducted in the presence of gangliosides. An increase in the hydrodynamic radius was observed for both A*β*40 and A*β*42, which suggests that the ganglioside lipids bind to the peptide monomers and form A*β*-ganglioside complexes. This is schematically represented in Figure 7a. Furthermore, studies have shown and hypothesised that GM1 gangliosides are in fact membrane binding sites for A*β*.^19, 40^ Additionally, simulation results have recently corroborated that GM1 gangliosides form complexes with both A*β*40 and A*β*42, and by binding monomers this process can result in the inhibition of the aggregation pathway.^28, 29^ In fact, the simulation results predicted that A*β*40 preferentially binds to GM1 gangliosides,^28^ which would explain why in our experimental results we observed complete inhibition of A*β*40 aggregation, while only partial inhibition of A*β*42. Therefore a potential mode of action is that ganglioside molecules create a complex with A*β* monomers and bind them in such a way that promotes a configuration whereby subsequent nucleation and fibrillar growth is impeded. The A*β* monomers are thus sequestered from the solution and the overall process of self-assembly is delayed. Such a mechanism could also be instrumental in how cells deal with extracellular A*β*. If gangliosides can form a complex structure with A*β* monomers, then cellular phagocytosis may be enhanced and A*β* can be cleared faster.^11^ If this mechanism is correct, this would also validate why ageing individuals, where ganglioside levels decline, ^21–24^ are more likely to develop neurodegenerative diseases.

**Figure 7:**
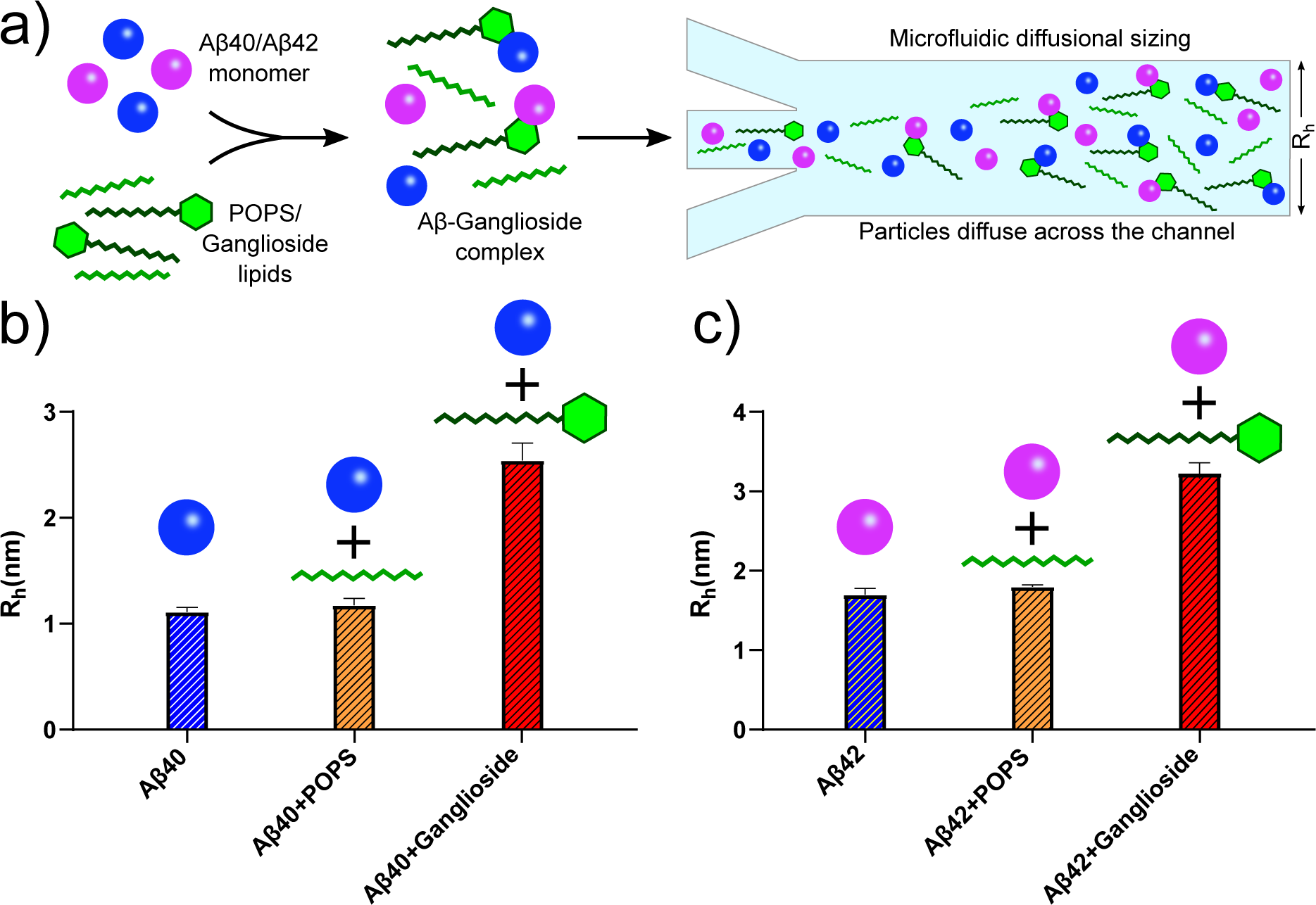
**(a)** Schematic representation of A*β*40/A*β*42 forming complexes with gangliosides but not with other lipids such as POPS. This was established using microfluidic diffusional sizing (MDS). **(b)** MDS results showing the hydrodynamic radii of A*β*40 monomers alone (blue bar), with POPS (yellow bar) and with ganglioside lipids (red bar). The change in hydrodynamic radius of A*β*40 with gangliosides but not with POPS suggests that the peptide formed a complex with gangliosides only. **(c)** Corresponding MDS results for A*β*42 under the same conditions, monomer alone (blue bar), with POPS (yellow bar) and with ganglioside (red bar). The data again suggest that a complex was formed with ganglioside lipids but not with POPS.

In conclusion, these findings further contribute to our understanding of how the aggregation of A*β* can be modulated by lipids which are predominantly found in the brain and how a reduction of certain lipids such as gangliosides can accelerate the primary nucleation rate and result in augmented formation of A*β* fibrils. Thus we shed light on the mechanistic processes behind some of the key initial steps underlying the aggregation process of A*β*42/A*β*40 in the presence of ganglioside lipids, that could be potentially used to inhibit the onset and proliferation of amyloid diseases such as Azlheimer’s Disease in the near future.

## Acknowledgements

We would like to acknowledge the EPSRC Underpinning Multi-User Equipment Call (EP/P030467/1) for funding the TEM in the Yusuf Hamied Department of Chemistry. Z.T. acknowledges funding from the Ron Thomson Research Fellowship, Pembroke College Cambridge. A.K.J acknowledges funding from the Cambridge Trust, the EPSRC grant EP/L015978/1 for the Centre for Doctoral Training for Nanoscience and Nanotechnology (NanoDTC), Queens’ College. T.P.J.K. acknowledges funding from the European Research Council under the European Unions Seventh Framework Programme (FP7/2007-2013) through the ERC grants PhysProt (agreement no. 337969), the Biotechnology and Biological Sciences Research Council (BBSRC), the Frances and Augustus Newman Foundation, and the Centre for Misfolding Diseases.

## Materials and Methods

### Aggregation kinetics

In order to probe and monitor the aggregation kinetics of both A*β*42 and A*β*40 with and without the presence of a ganglioside interface, a previously established protocol was followed^32^ A 96-well plate (Corning 3881, Half-area) was used. This has a cylindrical shape and thus the cross-sectional surface area remains the same throughout the well, making it ideal to monitor how the surface-to-volume (S/V) ratio, as well as the choice of interface, may affect protein aggregation. Moreover, it is important to note that the plates used in all experiments are specifically coated in order to ensure that peptides/proteins do not adsorb to the walls of the wells. Thus interactions from the walls of the wells were not considered in this study.

In order to monitor the aggregation process, 50 *µ*M Thioflavin T (ThT), a molecule that increases its fluoresce in the presence of *β*-sheets, was added to the solution. ThT fluorescence was monitored as a function of time using a microcplate reader (FLUOstar-BMG labtech). The temperature during all experiments was 37*^◦^*C. All experiments were performed under quiescent conditions without agitation.

For the experiments involving a change in the S/V ratio, different volumes of peptides were pipetted into the wells (80, 100, 120, and 140 *µ*L). This is schematically shown in Figure 2a-c.

### Amylofit kinetic analyis

All the kinetic data obtained was analysed using Amylofit.^41^ All data were initially normalised. The half-times were calculated from the time at which half the protein had aggregated, i.e. the time at which the normalised intensity reached 0.5. Each experiment was repeated three times and averaged before fitting. From the data, a secondary nucleation dominated model was found to be the best model to use, and thus this model was used throughout the study.

### Preparation of gangliosides, POPS and Purification of A***β***42 and A***β***40

A concentration of either 1 mg/mL Total Ganglioside Extract or 1mg/mL POPS was used (Avanti Polar Lipids). In brief, ganglioside/POPS lipids were dissolved in chloroform and then left to evaporate overnight in a fume hood. Following this, 1 mL of hexadecane (Merck) was added to the dried film and vortexed for 5 min to ensure complete mixing. The lipid solution was then added on top of the protein solution in each well of the 96-well plate, as shown in Figure 2a-c. The aggregation kinetics of A*β*42 and A*β*40 were then monitored and investigated as described above. The method of purifying A*β*42 and A*β*40 was taken from a previously established protocol.^42^ Following purification, the peptides were lyophilised and then redissolved in 20mM sodium phosphate buffer (pH 7.4 for A*β*42 and pH 8.0 A*β*40) at a stock concentration of 5 *µ*M (A*β*42) and 10 *µ*M (A*β*40).

### Seeded assay

For the experiments involving seeds, A*β*42 and A*β*40 were left to incubate (without the presence of a lipid interface) for one week at 37*^◦^*C. The aggregated peptides were then sonicated at 40% power for 30s which resulted in the formation of seeds. A 10% seed solution was then prepared for both A*β*42/A*β*40 and experiments were performed accordingly (ThT aggregation kinetics) as per the assays above. This is schematically shown in Figure 3a-c.

### Lipid mixtures

Lipid films consisting of POPC, POPS, POPE, POPG, PI4P, sphingomyelin, cholesterol and ceramides (Avanti Polar Lipids) were evaporated in a glass vial, in ratios mimicking the neuronal cell membrane as described in the lipidomics studies conducted by.^33^ To the resulting films, hexadecane (Merck) was added, and the experiments were conducted in a similar manner as described above.

### Transmission electron microscopy (TEM)

Transmission electron microscopy (TEM) was performed using a Thermo Scientific (FEI) Talos F200X G2 TEM operating at 200 kV. TEM images were acquired using a Ceta 16M CMOS camera. TEM grids (continuous carbon film on 300 mesh Cu) were glow discharged using a Quorum Technologies GloQube instrument at a current of 25mA for 60s. Samples were negatively stained using a 2% uranyl acetate solution for 45s.

### Cell culture of SH-SY5Y cells

SH-SY5Y cells (Merck), derived from the SK-N-SH neuroblastoma cell line, were cultured using Dulbecco’s Modified Eagle Medium (DMEM, Thermofisher Scientific), 10% foetal bovine serum (FBS, Merck) and GlutaMAX Supplement (Thermofisher Scientific). Following confluence, cells were seeded at a density of 10^5^ cells/cm2 in a 96 well plate. A*β*42/A*β*40 fibrils from the aggegration experiments were incubated with the cells overnight. All steps were performed in a sterile environment.

### Cytotoxicity and cell proliferation using MTT Assay on SH-SY5Y cells

The viability of SH-SY5Y cells following incubation with fibrils formed in the absence/presence of gangliosides was determined using a standard MTT cell proliferation assay (Merck).

In brief, 10 *µ*L of MTT reagent (3-[4,5-dimethylthiazol-2-yl]-2,5-diphenyltetrazolium bromide) labelling reagent Merck) was added to each well, followed by incubation for 4 hours. Subsequently, 100 *µ*L of solubilisation solution was added and the well plate was further placed in the incubator overnight at 37*^◦^*C and 5% CO_2_. The resulting absorbance of the solubilised formazan crystals was measured using a plate reader at 595 nm (FLUOstar Omega microplate reader BMG Labtech).

### Microfluidic Diffusional Sizing (MDS)

Microfluidic diffusional sizing (MDS) experiments were conducted using a previously established protocol.^43^ In brief, microfluidic PDMS devices (the master of which was fabricated using a soft lithographic process^44, 45^) were firstly coated with 0.01% Tween 20. Devices were then equilibrated for 1.5 minutes, before the experiment was run. Flow rates of 100-300 *µ*L/hr were used.

## Supporting Information

**Figure S1:**
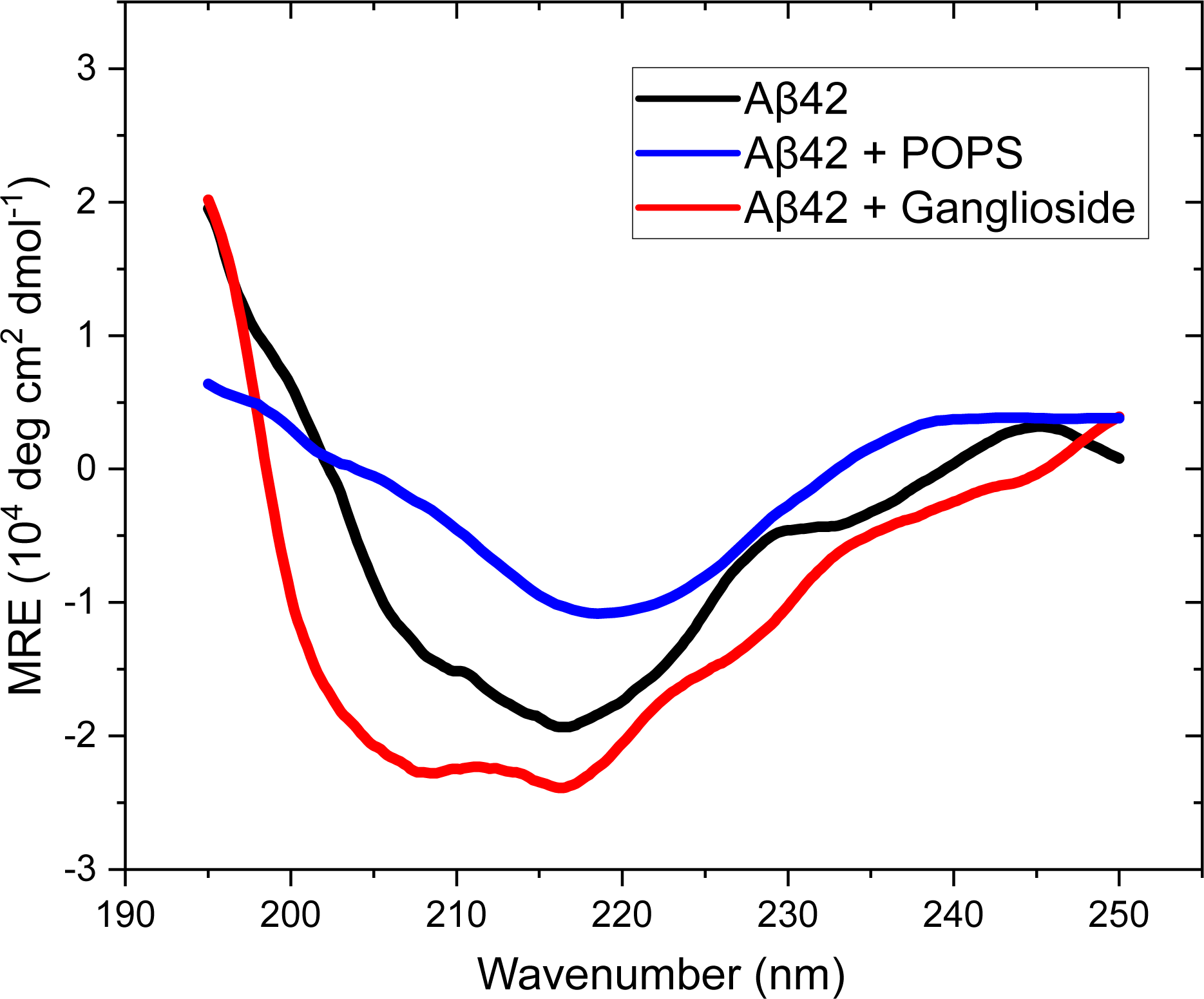
CD spectra of a 1 *µ*M A*β*42 solution following aggregation in the absence of a lipid (black curve) with POPS (blue curve) and with gangliosides (red curve). The negative band at 217/218nm is characteristic of a *β*-sheet structure.

**Figure S2:**
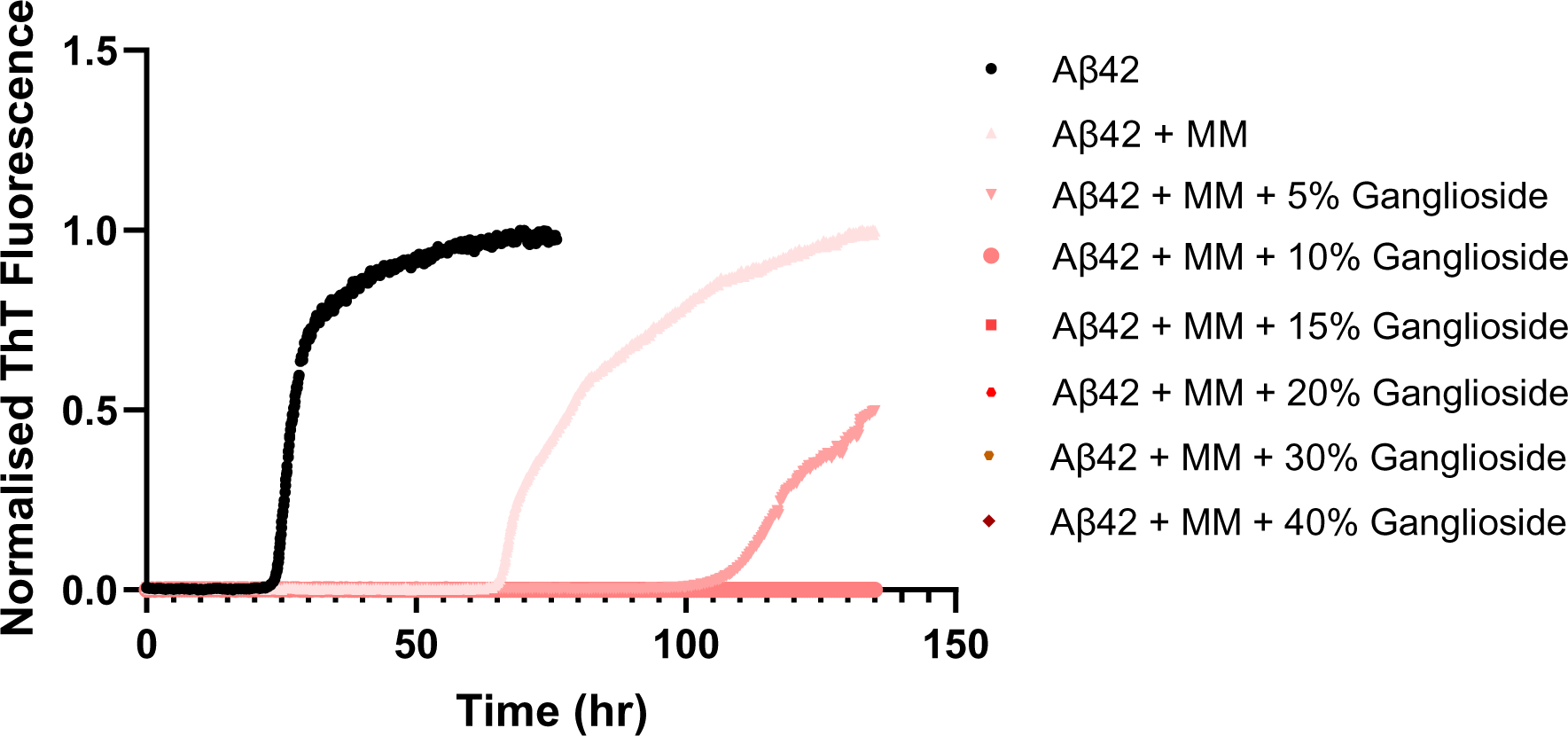
Normalised ThT aggregation kinetics of a 3 *µ*M A*β*40 solution in the absence and presence of a lipid mixture with varying amounts of gangliosides.

